# Evidence of persistent hyperphagia following a dietary weight-loss intervention in mice

**DOI:** 10.1101/2024.08.16.608348

**Authors:** Frankie D. Heyward, Evan D. Rosen

**Affiliations:** Division of Endocrinology, Diabetes, and Metabolism, Beth Israel Deaconess Medical Center; Harvard Medical School; Broad Institute of Harvard and MIT; Center for Hypothalamic Research, Department of Internal Medicine, UT Southwestern Medical Center, Dallas, TX, USA; Department of Neuroscience, UT Southwestern Medical Center, Dallas, TX, USA

## Abstract

**Objective:** This study sought to determine whether the drive to regain weight following weight loss was truly long-lived in mice.

**Methods:** We generated a model of reduced dietary obesity (ReDO) whereby male mice with diet-induced obesity (DIO) mice were calorically restricted until weight matched to control mice, and then after a 24-hour food assessment period were pair-fed relative to control mice. We subsequently generated ReDO mice that, after CR were pair-fed relative to control mice for 0, 8, or 28 days, or chronically. Body weight, food intake, and select metabolic parameters were measured, along with whole hypothalamic *Pomc* gene expression.

**Results:** ReDO mice in both experiments exhibited hyperphagia following CR, while a persistent form of hyperphagia was detected in ReDO_8d and ReDO_28d mice relative to control and chronically pair-fed mice. 4-week initial weight gain was predictive of the degree of weight regain across ReDO_8 and ReDO_28 mice.

**Conclusions:** ReDO mice exhibit a long-lived form of hyperphagia and an apparent drive to reclaim an upwardly shifted body weight set point. There was considerable variability with regard to ReDO_8 and ReDO_28 body weight regain which was correlated with the of initial degree of 4-week body-weight gain when first exposed to a high-fat diet. This study showcases the perdurance of weight loss-associated hyperphagia and introduces a prognostic tool for identifying mice that are prone towards weight regain, while setting the stage for future inquiries into the neurobiological basis of persistent hunger following weight loss owed to a dietary intervention in mice.

## INTRODUCTION

Obesity exerts a tremendous public health and economic burden within the US, therefore effective weight loss interventions are in high demand. Bariatric surgery has been an effective means of weight loss for many, yet almost 50%-80% of patients experience some degree of weight regain, with the percentage of regained weight ranging from 16 to 37%^1, 2, 3^. Additionally, new blockbuster drugs, such as the Glp1r agonist Semaglutide, have offered an unprecedented therapeutic benefit, with many losing as much as ∼24 of body weight^4^. Yet, there is evidence that 67% of the weight loss owed to Semaglutide returns within a year of halting the treatment, despite adherence to regimen^5^. Better understanding the physiological drivers of weight regain may give rise to therapeutic interventions that promote sustained weight loss.

Previous studies involving mouse models of diet-induced obesity (DIO) subjected to caloric restriction (CR), or a return to a chow diet, have observed that mice exhibit a long-lived obesogenic memory in the form of chronic inflammation in the periphery along with hyperphagia-driven weight regain ^6, 7, 8, 9^. Yet, to date, the documented assessments of weight regain-associated hyperphagia are initiated immediately following CR, making it unclear whether the heightened hunger exhibited by ReDO mice represents either an expected response to prolonged exposure to caloric restriction, or a drive to re-establish a long-lived upwardly shifted (relative to control mice) homeostatic body weight set point. Thus, we set out to determine the longevity of weight-loss-associated hyperphagia in mice.

## METHODS

### Animals

All animal experiments were performed with approval from the Institutional Animal Care and Use Committees of The Harvard Center for Comparative Medicine and Beth Israel Deaconess Medical Center. 4-week-old (experiment 1) or 5-week old (experiment 2) C57BL/6J male mice from The Jackson Laboratory (Strain #000664) were shipped to our animal facility in groups of weanlings. Upon arrival, mice were group-housed (*n* = 5), with food and water available *ad libitum*, on a 12:12 h light/dark schedule [lights on at 6 A.M (experiment 1) or 7 A.M (experiment 2).; zeitgeber time (ZT) 0]. Mice were acclimated to the housing conditions for at least 1 week. At 6 weeks of age, mice were maintained on either a control, standard chow, diet or a DIO (60% fat by calories from lard) diet (Research Diets, #D12492) for 16 weeks (experiment 1) or 20 weeks (experiment 2). Individual mouse body weights were measured weekly prior to the CR phase of the experiment. One control mouse’s 24-hour food intake measurement was omitted owed to it being recorded as 153 g, which is physically impossible (experiment 1).

### Measurement of endocrine profiles

Fasting glucose levels were obtained by fasting mice overnight for 12 hours, after which tail blood glucose levels were measured using a glucometer. Serum samples for insulin were taken from trunk blood following euthanasia and decapitation. Insulin analysis was performed via ultra-sensitive mouse insulin ELISA kit (Crystal Chem, Downers Grove, IL).

### Caloric restriction and paired feeding

Experiment #1: Prior to CR all mice were group-housed. Control mouse cage-wide food intake was determined for 24 hours, and group-housed CR mice were given that amount the following day to commence CR. After 1 week (week 17) all mice were singly housed and ReDO mouse daily food intake was measured across two days, after which mice were given 60% of this amount for 12 days until the CR phase was complete. Immediately following CR food intake was measured for 24 hours for all mice, prior to initiation of paired feeding. Mice were weighed daily and given food 30 minutes to 2 hours prior to the onset of the dark cycle.

Experiment #2: Control, DIO, and ReDO mice were singly housed after week 20. Control mouse 24-hour food intake was measured across two days to determine the two-day average food take, then 60% of this food weight was calculated to determine the amount of food to administer to achieve 40% caloric restriction (e.g., 3.45g x 0.60 = 2.1g).

### Real-Time PCR Analysis

Whole hypothalamus was collected in Trizol reagent (Thermo Fisher), RNA was isolated, cDNA generated and qRT-PCR was performed with normalization involving the housekeeping gene *Hprt*. Analysis of qPCR data was conducted via the 2^-ΔΔCT^ method.

### Quantification and statistical analysis

Data are expressed as means ± SEM. Analyses were performed using GraphPad Prism (GraphPad Software, San Diego, USA). Data were compared between pairs of groups using the Student t-test, or One-way ANOVA with Holm-Sidak multiple comparison test, and correlation analyses were performed by Pearson correlation, owed to the data being normally distributed.

## RESULTS

We first sought to establish a successful model of diet-induced obesity (DIO) followed by reduced dietary obesity (ReDO) in mice. To this end, we employed both male control mice and those subjected to diet-induced obesity (DIO) fed a high-fat diet (60% fat), with both groups maintained on their respective diets for 16 weeks. DIO mice weighed substantially more than control mice and exhibited hyperglycemia (**Figure 1a,b,d**). DIO mice were then fed either a 40% caloric restriction diet (ReDO) until their aggregate body weight was matched to that of control mice, which occurred after 19 days of CR. Mice were then allowed to consume chow diet *ab libitum*, in an attempt to stabilize their weights. Interestingly, ReDO mice exhibited a high degree of hyperphagia during a 24-hour period (**Figure 1c**). To match their weight to that of control mice, ReDO mice were pair-fed with their control counterparts for an additional 7 days. While percent body fat, as measured by Dual-energy X-ray absorptiometry (DEXA), was comparable between control mice and ReDO mice (**Figure 1e**), and ReDO adipocyte diameter appeared similar as well, ReDO mice exhibited a pronounced degree of adipocyte inflammation as indicated by the existence of crown-like structures around in their epididymal fat adipocytes (**Figure 1f**).

**Figure 1.**
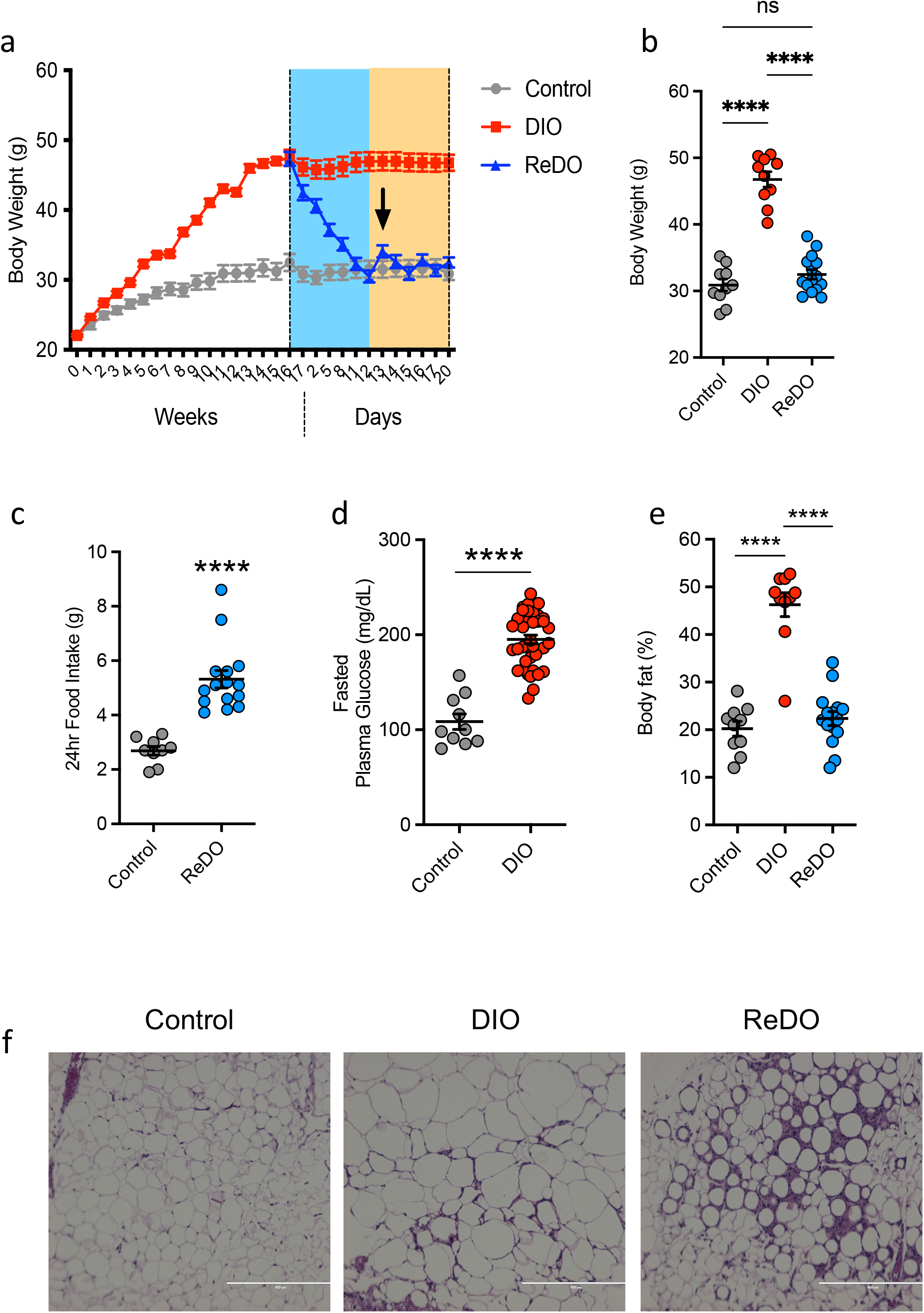
ReDO mice exhibit increased hunger immediately following CR. (**a**) Weekly body weight measurements 16 weeks followed by daily body daily body weight measurements during the CR phase and beyond. (**b**) Body weight measured at the conclusion of CR. (**c**) 24 hour food intake comparing control and ReDO mice. (**d**) Fasted glucose. (**e**) % Body Fat. (**f**) Representative histology images across control, DIO, and ReDO groups.

Intrigued by the marked degree of hyperphagia displayed by ReDO mice, we next sought to determine if the drive to consume food was owed to the mice having immediately switched from a hunger-inducing state to a post-CR *ab libitum* state, as opposed to a response that was driven by some enduring orexigenic drive despite being temporarily removed from their exposure to CR. To test this, control or DIO were maintained on their respective diets for 20 weeks, at which time mice were subjected to 40% CR for 3 weeks until they were weight-matched to their lean control counterparts (**Figure 2a, b**). Next, ReDO mice were divided into 4 distinct groups, ReDO_0d, ReDO_8d, ReDO_28d, or ReDO_pf, which were pair-fed relative to control mice, for 0 days, 8 days, 28 days, or perpetually, respectively (**Figure 2a**). All ReDO groups, with the exception ReDO_pf, were allowed to consume food *ad-libitum* at the end of their paired-feeding regimen.

**Figure 2.**
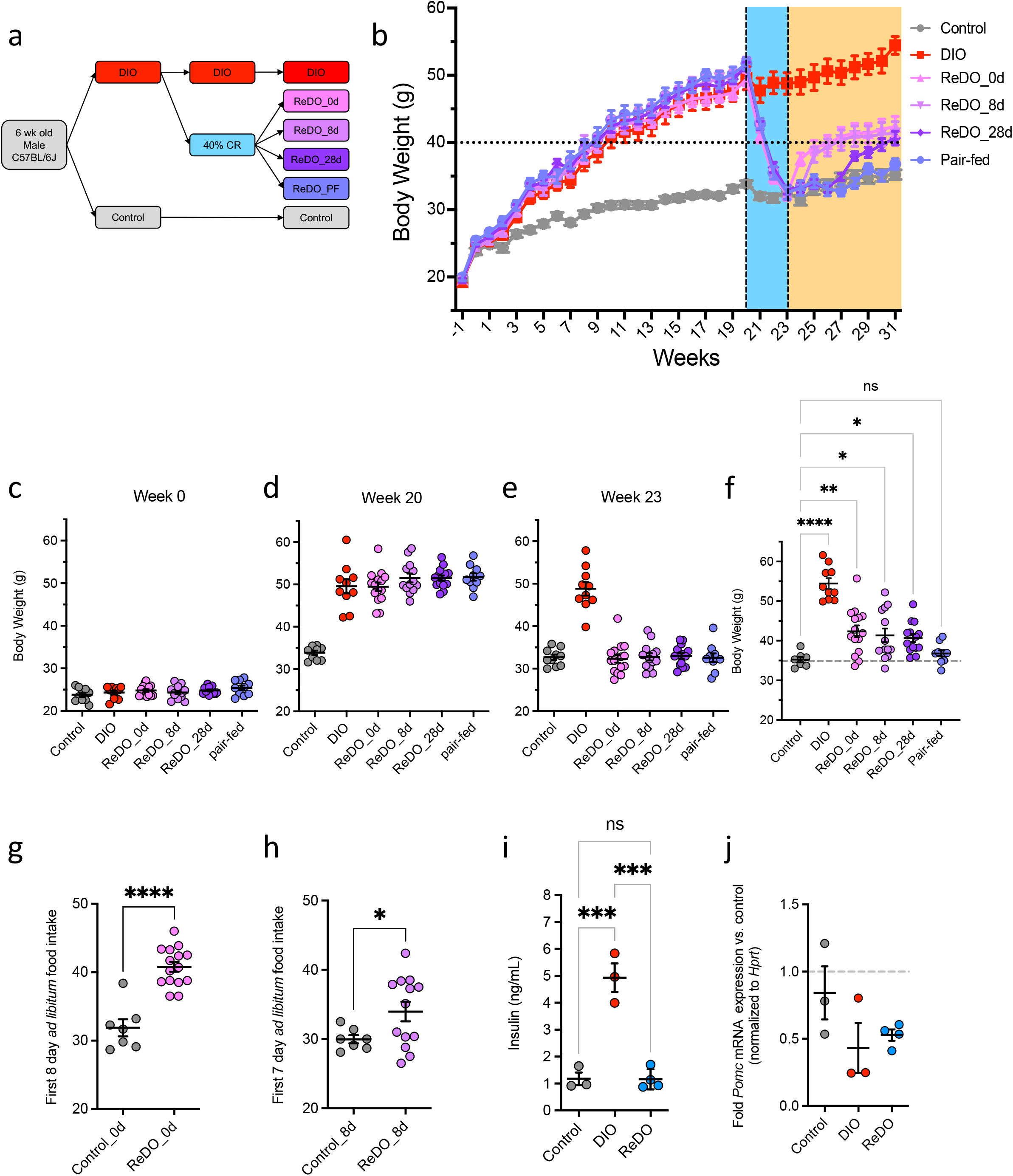
ReDO mice exhibit a persistent hyperphagia and weight gain phenotype. (**a**) Schematic of the experimental design. Male control mice were fed a standard chow diet, while DIO mice were maintained on a high-fat diet (HFD, 60% kcal from lard), for 20 weeks prior to 40% caloric restriction until reduced dietary obesity (ReDO) mouse groups were weight matched to lean control mice. ReDO mice were pair-fed using a standard chow diet for either 0 days, 8 days, or 28 days, prior to being maintained on an *ad libitum* chow diet, while ReDO_pf mice were paired fed chronically. (**b**) Body weight across the experiment depicting the control, DIO, ReDO_0d, ReDO_8d, ReDO_28d, and ReDO_pf groups. (c) Body weight at week 0. (d) body weight at week 20. (e) Body weight at week 23. (f) Body weight at week 31. (g) First 8-day *ad libitum* food intake for ReDO_0d mice. (h) First 7-day *ad libitum* food intake for ReDO_8d mice. (i) Fasted serum insulin. (j) *Pomc* gene expression.

Mice were pre-assigned to their respective groups at the onset of the experiment, ensuring that all groups started with average weights that were not significantly different (**Figure 2c**). After having been maintained on their diet for 20 weeks, all DIO groups, including future ReDO groups, weighed significantly more than control mice while not differing amongst themselves (**Figure 2b, d**). 7 mice appeared to be diet-induced obesity resistant (DIO-R), according to the criteria used in Enriori et al., (2007), as they possessed a body weight that was within ± 3 standard deviations of the average body weight on control mice, and were therefore removed from the experiment.

Despite being weight matched to control mice after CR, all ReDO groups, regardless of the duration of paired feeding, weighed significantly more than control mice 4 weeks after re-exposure to a standard chow *ad libitum* diet (**Figure 2b, f**). Importantly, ReDO_pf mice did not exhibit an appreciable degree of weight gain, supporting the conclusion that the weight regain in our ReDO model was principally driven by persistent hyperphagia. Indeed, both ReDO_0d and ReDO_8d mice displayed elevated cumulative food intake, 8 days and 7 days, respectively, after the resumption of *ad libitum* chow-diet feeding. Food intake data for the ReDO_28d 1-week *ad libitum* feeding reintroduction are not reported due to a scale malfunction, which precluded accurate measurement.

To ascertain the physiological basis for the ReDO mouse weight regain, we sought to determine if these mice, despite being weight-matched to controls, were metabolically dysfunctional. So as to not confound our experiment, we measured fasted serum insulin levels in a subset of ReDO_8d mice immediately prior to *ad libitum* feeding, which required their euthanasia and removal from the remainder of the experiment. ReDO serum insulin levels were indistinguishable from that of controls (**Figure 2i**). In these same mice, we measured *Pomc* mRNA expression in whole-hypothalamus, which trended down in comparison to that of control mice but was non-significant (**Figure 2j**).

Given the considerable degree of regained body weight stratification within each of our ReDO groups, we sought to identify parameters that could be used to predict the degree of weight regain during future ReDO experiments. Correlational matrixes were generated for ReDO_0d, ReDO_8d, and ReDO_28d mice, comparing up to 12 variables (ReDO_28 mice are missing two food-intake-related variables) (**Figure 3a,d,g**). Given the Gaussian distribution of our data, we generated Pearson Correlation Coefficients and p-values to assess the strength of the linear relationship between two variables. Interestingly, for both pair-fed groups ReDO_8d and ReDO_28d, initial 4-week body weight (BW) change upon initial exposure to a high-fat diet was strongly correlated with the ultimate degree of body weight regain (**Figure 3d,f,g,h**).

**Figure 3.**
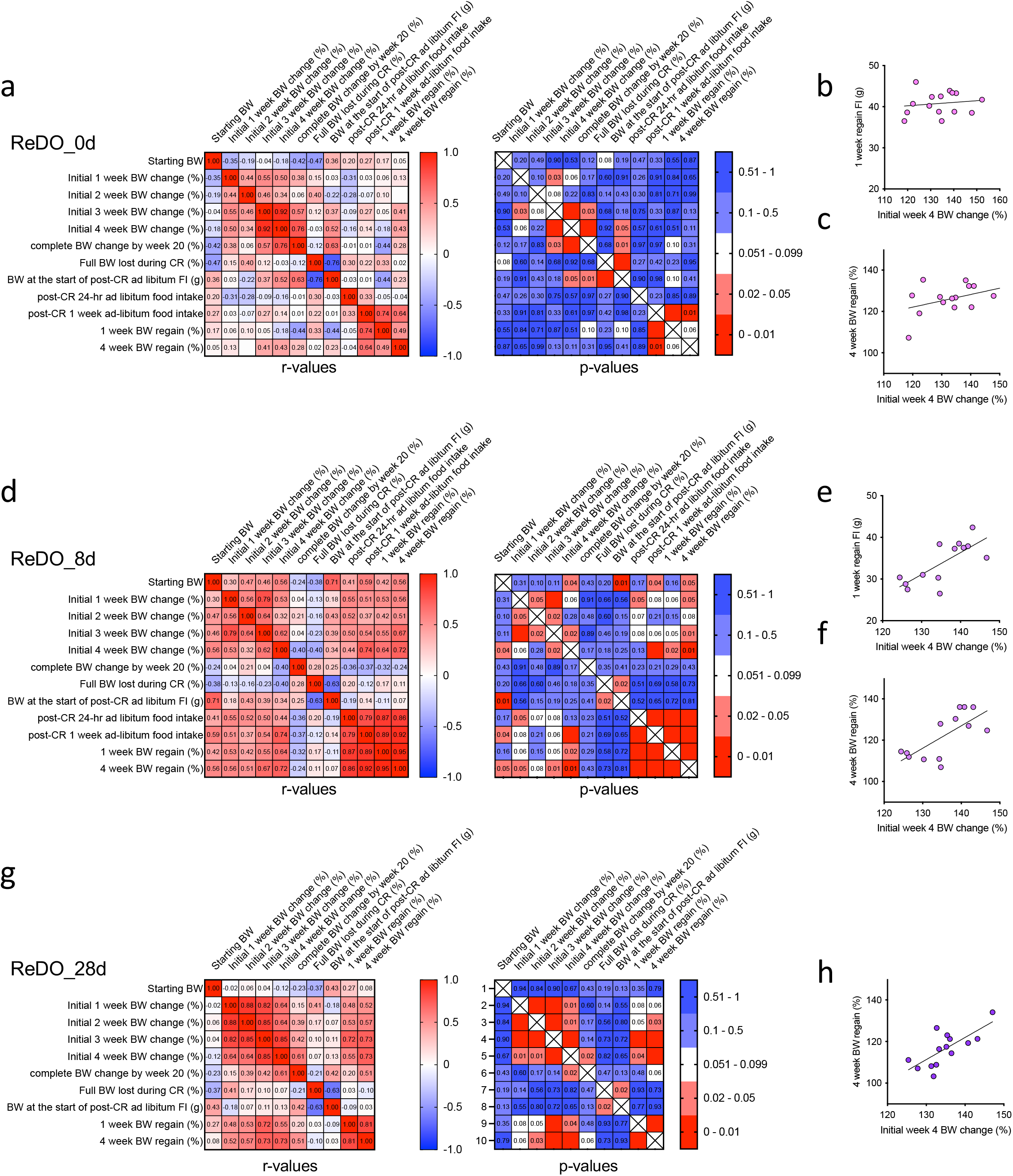
Initial body weight change is correlated with the degree of weight regain. (a) Heatmaps of Pearson correlation r values (left) and p values (right) between various parameters for ReDO_0d mice. (b) Linear regression comparing ReDO_0d Initial week 4 BW change versus 1 week regain food intake. (c) Linear regression comparing ReDO_0d Initial week 4 BW change versus 4-week BW regain. (d) Heatmaps of Pearson correlation r values (left) and p values (right) between various parameters for ReDO_8d mice. (e) Linear regression comparing ReDO_8d Initial week 4 BW change versus 1 week regain food intake (g). (f) Linear regression comparing ReDO_8d Initial week 4 BW change versus 4-week BW regain. (g) Heatmaps of Pearson correlation r values (left) and p values (right) between various parameters for ReDO_28d mice. (h) Linear regression comparing ReDO_28d Initial week 4 BW change versus 4-week BW regain. Food intake columns are missing for ReDO_28d mouse comparisons.

## DISCUSSION

Our findings recapitulate and extend those of other groups, revealing that mice with reduced dietary obesity are not only hyperphagic immediately following a CR regimen ^6^, but their hyperphagia lasts for at least a month, despite being diet-matched to control mice during this time. Whereas previous studies have detected various forms of obesogenic memory in the periphery (e.g., persistent adipose tissue inflammation)^7^, it’s unclear whether these contribute to weight regain. Attention should be directed toward identifying the key drivers of weight-loss-associated hyperphagia. Various neuronal populations within hypothalamus, hindbrain, and other brain regions have been established as key regulators of hunger^10^. In particular, leptin receptor-expressing neuronal-types within the hypothalamus, including AgRP and Pomc neurons, as well as Glp1r-expressing cell-types within the hindbrain and hypothalamus, have been strongly implicated in the control of hunger and body weight ^11, 12, 13, 14, 15^. Our study, similar to that of earlier work^6^, failed to detect significant hypothalamic gene-expression profiles indicative of impairments in the function of these neuronal systems, likely owed to the technical difficulties of measuring cell type-intrinsic gene expression profiles in “noisy” whole-tissue preps with an exceptionally high degree of cellular heterogeneity. Future studies should infer the state of hunger-linked neurons in ReDO mice using sensitive cell-type specific transcriptomic approaches optimized for rare hypothalamic neuronal types ^16^.

Compared to lean control mice, mice with reduced dietary obesity appear to be driven to reclaim an upwardly shifted body weight set point. Precisely when, after initial exposure to a high-fat diet, this set point becomes upwardly shifted remains an open question. Recent work involving an intragastric overfeeding mouse model suggests that after 14 days mice rapidly re-establish a weight that is similar to control mice suggesting set point shifting occurs after more than 2 weeks of sustained positive energy balance^17^. Moreover, another study, using DIO mice that were switched back to chow in a staggered fashion, suggests that set point shifting may occur between 8 weeks and 24 weeks of HFD exposure^18^.

We observed a fair degree of population structure within our data, with some mice regaining more weight, and consuming more food, than others. Given the low degree of body weight variability within each of the ReDO groups prior to HFD feeding, we suspect that some aspect of high-fat diet exposure unveils a hidden susceptibility to diet-induced obesity. Furthermore, 4-week body weight change being positively correlated with weight regain in ReDO_8d and ReDO_28d mice suggests that mice may be differentially programmed for a unique susceptibility to weight regain that is distinct from a general susceptibility to initial weight gain. Indeed, total % weight gain, prior to CR, was not correlated with weight regain. We suspect ReDO_0d do not show this association because their post-CR orexigenic drive is so uniformly strong that it masks any within-group differences. The initial 4-week body weight change metric can be leveraged to identify weight-regain susceptible versus resistant mice *a priori* for future comparative studies.

## Acknowledgments

This work was supported by an NIH 3 T32 DK 7516-32 to F.D.H., and R01 DK085171 to E.D.R.

## Author Contributions Statement

F.D.H. conceived this study, conducted all experiments, and interpreted the results of all experiments. E.D.R supervised this study and provided funding. F.D.H wrote the manuscript with added input from E.D.R.

## Competing Interests Statement

The authors have no competing interests to disclose.

## Notes

### Competing Interest Statement

The authors have declared no competing interest.

